# ATRIP protects progenitor cells against DNA damage *in vivo*

**DOI:** 10.1101/2020.02.14.948265

**Authors:** Gabriel E. Matos-Rodrigues, Paulius Grigaravicius, Bernard S. Lopez, Thomas Hofmann, Pierre-Olivier Frappart, Rodrigo A. P. Martins

## Abstract

The maintenance of genomic stability during the cell cycle of progenitor cells is essential for the faithful transmission of genetic information. Mutations in genes that ensure genome stability lead to human developmental syndromes. Mutations in Ataxia Telangiectasia and Rad3-related (ATR) or in ATR-interacting protein (*ATRIP*) lead to Seckel syndrome, which is characterized by developmental malformations and short life expectancy. While the roles of ATR in replicative stress response and chromosomal segregation are well established, it is unknown how ATRIP contributes to maintaining genomic stability in progenitor cells *in vivo*. Here, we generated the first mouse model to investigate ATRIP function. Conditional inactivation of *Atrip* in progenitor cells of the CNS and eye led to microcephaly, microphthalmia and postnatal lethality. To understand the mechanisms underlying these malformations, we used lens progenitor cells as a model and found that ATRIP loss promotes replicative stress and TP53-dependent cell death. *Trp53* inactivation in *Atrip*-deficient progenitor cells rescued apoptosis but increased mitotic DNA damage and mitotic defects. Our findings demonstrate an essential role of ATRIP in preventing DNA damage accumulation during unchallenged replication.

## Introduction

Genomic stability is crucial for cellular and tissue homeostasis, normal development, and the prevention of tumorigenesis and aging. Mutations in genes that ensure genome stability and proper chromosomal segregation lead to human developmental syndromes ^1, 2^. DNA damage response (DDR) pathways sense, signal and repair different types of DNA damage and are crucial for maintaining genomic stability ^3^. The Ataxia Telangiectasia and Rad3-related (ATR) kinase regulates replication fork firing and stability to ensure the proper timing of DNA replication during S-phase and cell cycle checkpoints to prevent mitosis in cells with underreplicated DNA ^4, 5^. When single-stranded DNA (ssDNA) is exposed at DNA damage sites or replication forks, RPA-coated ssDNA recruits ATR-interacting protein (ATRIP) and activates ATR, which coordinates a replicative stress response (RSR) ^6, 7^. ATR protein stability and activity as well as all the described functions of ATR depend on ATRIP ^6, 8^.

In humans, mutations in *ATR* or *ATRIP* lead to Seckel syndrome, which is characterized by growth defects, neurodevelopmental malformations and short life expectancy ^1^. Mouse models of ATR loss of function have highlighted the essential contribution of ATR to development. While *Atr* germline inactivation leads to early embryonic lethality ^9, 10^, conditional knockout mice have revealed that ATR plays essential roles in proper cell cycle progression, genome stability and meiosis ^11, 12, 13, 14^. In addition, hypomorphic Atr mutations in mice recapitulate some of the developmental defects observed in Seckel syndrome patients ^15^. Although the contribution of ATR to genomic stability and appropriate development has been extensively investigated, the function of ATRIP *in vivo* is unknown.

The accumulation of DNA damage can activate the tumor suppressor protein TP53, a master regulator of the DDR that regulates cell cycle arrest, apoptosis, cell metabolism and DNA repair ^16, 17^. Moreover, inactivation of DDR-related genes may lead to TP53-dependent apoptosis ^17, 18^. Notably, *Atr* inactivation increases spontaneous DNA damage, but the simultaneous inactivation of Atr and Trp53 in mouse neural progenitor cells (NPCs) does not rescue brain growth defects ^12, 19^. Additionally, *TP53*-inactivated cancer cells are particularly sensitive to ATR signaling inhibition ^20, 21, 22^, suggesting that ATR and TP53 work in parallel to prevent DNA damage accumulation. Indeed, *Trp53* inactivation in hypomorphic Atr-mutated mice leads to synthetic lethality ^15^. This evidence suggests cooperation between the ATRIP-ATR signaling pathway and *Trp53*; however, the synergy between these pathways during the cell cycle progression of unchallenged progenitor cells is not completely understood.

Here, we developed a new mouse model to investigate the function of ATRIP. We showed that the tissue-specific inactivation of *Atrip* during CNS and eye development led to severe growth defects that were associated with the death of progenitor cells. Next, due to its well-characterized developmental kinetics and cell cycle dynamics ^23^, we used the developing lens as a model to better understand the cellular and molecular outcomes of *Atrip* inactivation *in vivo*. Importantly, the developing lens was particularly useful for probing the cross-talk between *Trp53* and other tumor suppressor genes ^24^. *Atrip* inactivation led to replicative stress, DNA damage accumulation and TP53-dependent apoptosis. Interestingly, while *Trp53* inactivation rescued the apoptosis of Atrip-deficient lens progenitor cells, it increased mitotic defects and did not rescue the observed microphthalmia. Taken together, these results reveal essential roles of ATRIP for DNA damage avoidance and progenitor cell homeostasis during development.

## Materials and Methods

### Mice

The experimental procedures involving animals were approved by the Committee of Ethics in Animal Use of the Health Science Center (CEUA/CCS) of Brazil and approved by the governmental review board of Baden-Württemberg, Germany (Regierungspräsidium Karlsruhe-Abteilung 3-Landwirtschaft, Ländlicher Raum, Veterinär-und Lebensmittelwesen).

The generation of *Atrip*^Flox^ mice was performed as follows. The targeting vector was constructed using the recombineering method developed by the National Cancer Institute (USA). The BAC containing *Atrip* was obtained from the Wellcome Trust Sanger Institute (bMQ-176B24). Exons 1, 2 and 3 of *Atrip* were targeted since the deletion of these exons was predicted to abrogate protein translation. One LoxP site was inserted upstream of exon 1, and the pGK-neo cassette flanked by two Frt sites followed by a second LoxP was inserted between exons 3 and 4. Prior to E14.1 ES cell electroporation, the targeting vector was linearized (NotI), and proper targeting was verified by restriction mapping and sequencing. After 12 days, 384 individual clones were analyzed by Southern blotting (with EcoRI digestion and a genomic probe), and 15 clones (3.9%) presented the expected targeting event. The presence of the third LoxP site was confirmed by PCR in 6 clones (Figure 1). The pGK-neo cassette was excised *in vivo* to generate the *Atrip* floxed allele using the FLPe mouse.

**Figure 1.**
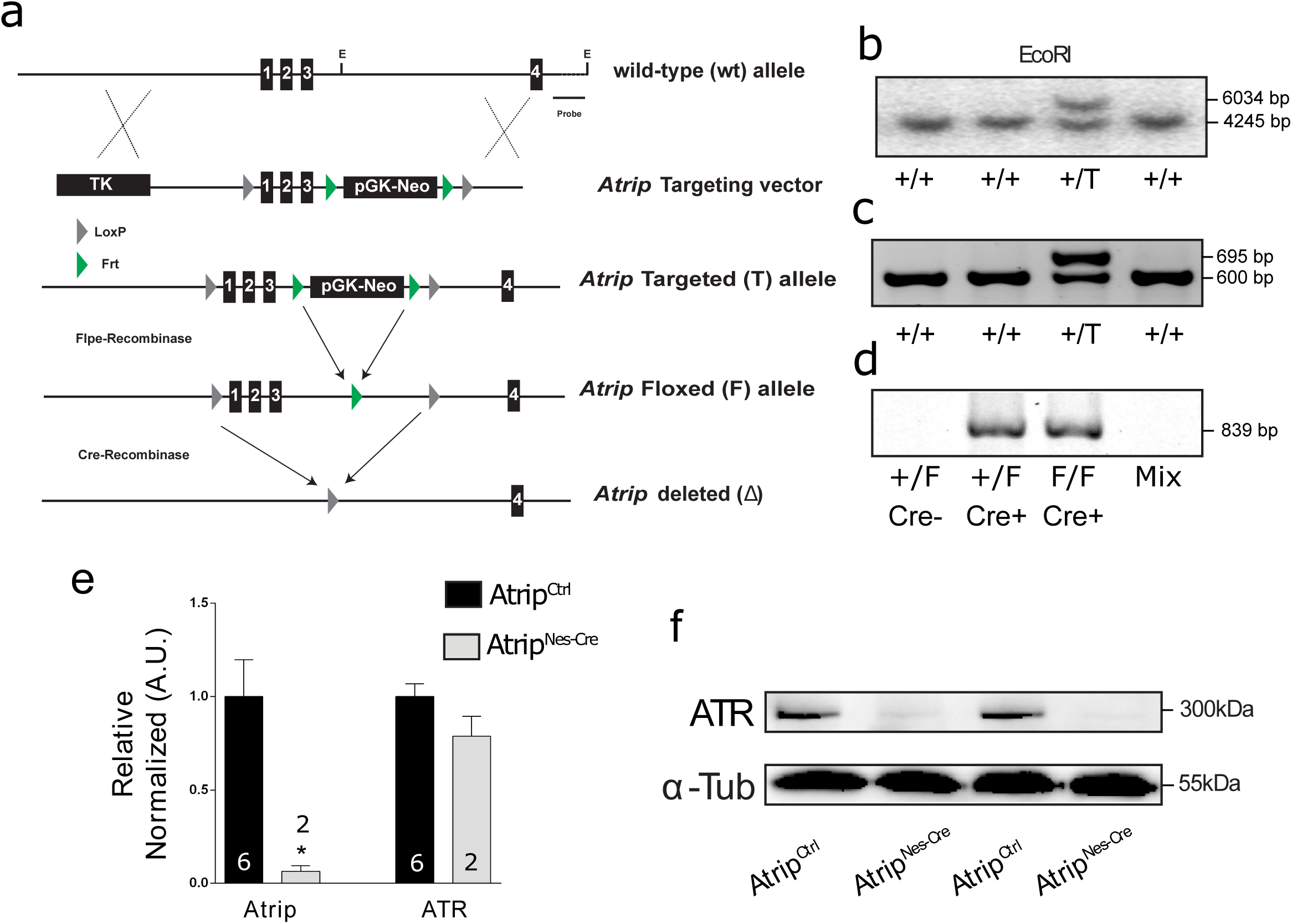
Generation of *Atrip* conditional knockout mice: **(a)** Gene-targeting strategy for the *Atrip* gene. The LoxP sequences flanking the three first exons of *Atrip* enable its genetic inactivation following Cre-mediated recombination. The pGK-Neo region of the transgene was excised *in vivo* using FLPe recombinase. **(b)** Southern blot generated using a genomic probe targeting intron 4 and digestion with EcoRI. **(c)** PCR amplification of the third LoxP site located upstream of exon 1. **(d)** PCR analysis of the Cre-mediated recombination of the *Atrip*^*flox*^ transgene in *Atrip*^Nes-Cre^ cortex. **(e)** *Atrip* and *ATR* mRNA expression analysis by real-time RT-PCR using cortex samples from *Atrip*^*Ctrl*^ and *Atrip*^*Nes-Cre*^ mice at E17,5. **(f)** Western blot of independent *Atrip*^*Ctrl*^ and *Atrip*^*Nes-Cre*^ cortex samples at E17,5. The number of biological samples analyzed is indicated in the bars. Error bars indicate SEM; *p<0.05.

*Nestin-Cre* (B6. Cg-Tg(Nes-cre)1Kln/J), *Lens-Cre* (Tg(Pax6-cre,GFP)1Pgr), *Trp53* floxed (FVB.129-Trp53tm1Brn) and FLPe (B6;SJL-Tg(ACTFLPe)9205Dym/J) mice were purchased from Jackson Laboratory. The control group (*Atrip*^*Ctrl*^) was composed of *Atrip*^*+/+*^, *Atrip*^*+/Flox*^ and *Atrip*^*Flox/Flox*^ mice. Mice in which *Atrip* was homozygously inactivated using *Nestin-Cre* were identified as *Atrip*^*Nes-Cre*^ = *Atrip*^*Flox/Flox*^; *Nestin-Cre*^*+/-*^. Mice in which *Atrip* and *Trp53* were homozygously inactivated using Nestin-Cre were identified as *Atrip;Trp53*^*Nes-Cre*^ = *Atrip*^*Flox/Flox*^; *Trp53*^*Flox/Flox*^; *Nestin-Cre*^*+/-*^. Mice in which *Trp53* was homozygously inactivated using *Nestin-Cre* were identified as *Trp53*^*Nes-Cre*^ = *Trp53*^*Flox/Flox*^; *Nestin-Cre*^*+/-*^. Mice in which *Atrip* was homozygously inactivated specifically in the lens were identified as *Atrip*^*Le-Cre*^ = *Atrip*^*Flox/Flox*^; *Le-Cre*^*+/-*^. Mice heterozygous for *Atrip* in the lens were identified as *Atrip*^*Het*^ = *Atrip*^*+/Flox*^; *Le-Cre*^*+/-*^.

### RNA extraction, cDNA synthesis and real-time RT-PCR

Lenses from three different mice of the same litter were dissected in cold PBS and lysed in 1 mL of TRIzol (Life/Thermo Fisher, Cat# 15596026), and RNA extraction was performed as previously described ^25^.

Real-time RT-PCR was performed in 96-well optic plates (Applied Biosystems, N801-0560) in an Applied Biosystems ABI7500 thermocycler. The following primers were used for real-time RT-PCR: SYBR primers, ATR forward 5’-TGCTATTCAGGAGTTGCTTTCT-3’ and reverse 5’-GGACATGCTCAGGGAATCTTT-3’ and Atrip forward 5’-TCTCCAGAAAGCTCCAATCAC-3’; TaqMan primers and probe, GPI1 forward 5’-TCCGTGTCCCTTCTCACCAT-3’, reverse 5’-GGCAGTTCCAGACCAGCTTCT-3’, and probe 5’-CTCCCTGCCCAGAGCGCACC-3’ and β-actin forward 5’-AGCCACCCCCACTCCTAAGA-3’, reverse 5’-TAATTTACACAGAAGCAATGCTGTCA-3’, and probe 5’-ATGGTCGCGTCCATGCCCTGA-3’. Each sample was run in duplicate, and only duplicates showing a variation of less than 0.5 Ct were further analyzed. The comparative Delta-Delta Ct (2-ΔΔCt) method for relative quantification was applied to determine the relative quantity of a target compared to the average of the two reference genes (GPI1 and β-actin). We employed a mathematical correction similar to QBASE software based on the use of the mean ΔCt of all groups to define the calibrator (Hellemans et al., 2007).

### Volume measurements of the eye

The volumes of postnatal and adult eyes were measured as previously described ^26^.

### Histology and immunofluorescence

Histological analyses were performed in cryosections. Embryo heads were fixed in 4% PBS-buffered paraformaldehyde for 24 hours at 4 °C and cryoprotected in 10, 20 and 30% (overnight) PBS-buffered sucrose solutions. Sections of 10 μm were obtained using a Leica 1850 cryostat.

Slides were washed with PBS, and antigen retrieval was performed using citrate buffer (pH=6). The following antibodies and dilutions were used: anti-Ser10 pH3 (1:50, Cell Signaling, cat# 9701), anti-active caspase-3 (1:100, BD Biosciences, cat#: 559565), anti-γH2AX (1:300, Millipore, cat# 05-636), anti-γH2AX (1:200, Abcam, cat# ab11174), anti-BrdU (1:3, General Electric, cat# RPN20), anti-Trp53 (CM5, 1:300, Vector Laboratories, cat# VP-P956), and anti-phospho TP53 (Ser15) (1:100, Cell Signaling Technology, #9284S).

Immunofluorescence reactions were developed via different methods, involving the use of a biotinylated secondary antibody followed by an ABC complex and Cy3-tyramide kit (Perkin Elmer, cat#: FP1046) or an Alexa secondary antibody (1:500, Invitrogen). Fluorescent nuclear counterstaining was performed using DAPI (Lonza, cat#: PA3013).

To label S-phase cells *in vivo*, intraperitoneal injections of 50 μg/g of body weight of BrdU (Sigma Aldrich, cat# B5002) were performed. Embryos were collected 1 hour after injection. TUNEL analysis was performed following the manufacturer’s instructions (Promega, cat# G7362). Fluorescent images were captured using a Leica TCS-SP5 with an AOBS system.

### Western blot analysis

Protein extraction and SDS-PAGE protein separation were performed as previously reported ^27^. Primary antibodies for the following target proteins were used: ATR (1:500, Santa Cruz Biotechnology, cat# SC1887) and α-tubulin (1:10000, Santa Cruz, cat #: sc32293). HRP-conjugated secondary antibodies from Cell Signaling were employed (1:1000, anti-mouse IgG, cat #: 7076, anti-rabbit IgG, cat #: 7074). The ECL system (cat #: RPN2132) was used according to the manufacturer’s instructions, and chemiluminescence was captured using ChemiDoc MP (BioRad) equipment.

### Statistical analysis

GraphPad Prism software was used for statistical analysis. The t-test, one-way ANOVA and two-way ANOVA were performed as indicated. The reported p-values are based on two-sided tests.

## Results

### ATRIP loss in the eye and CNS causes microcephaly, microphthalmia and premature lethality

To understand the function of ATRIP *in vivo*, we generated an *Atrip* conditional knockout mouse in which the first three exons of the *Atrip* gene were flanked by LoxP sequences (*Atrip*^*Lox*^) (Figure 1a). Three *Atrip*-targeted ES cell clones verified by Southern blotting were injected into blastocysts, and 12 chimeras with a high percentage of chimerism were obtained. Germline transmission was confirmed by Southern blotting and PCR (Figure 1b and c). To inactivate *Atrip* in the developing CNS and eye, we generated *Nestin-Cre; Atrip*^*Lox/Lox*^ mice (*Atrip*^*Nes-Cre*^). We confirmed genomic recombination by PCR (Figure 1d) and decreased *Atrip* mRNA expression by real-time RT-PCR (Figure 1e). Interestingly, no difference in *Atr* mRNA levels was detected in the cortex of *Atrip*^*Nes-Cre*^ mice at E17.5 (Figure 1e), but a decrease in the ATR protein content was evident (Figure 1f). These data indicate that, as previously described in cell lines ^8, 28^, ATRIP is essential for ATR protein stability in NPCs *in vivo*.

The inactivation of *Atrip* in the CNS and eye did not change the Mendelian ratio at birth in the mice; however, *Atrip*^*Nes-Cre*^ mice showed a reduced body mass (Figure 2a) and died around P9. Additionally, *Atrip*^*Nes-Cre*^ mice exhibited microcephaly and microphthalmia (Figure 2b and d). Histological analysis of P7 brains revealed profound developmental growth impairment of the cerebellum, hippocampus and dorsal striatum (Figure 2c). Notably, these phenotypes mirror the developmental defects observed in mice with an ATR-deficient CNS ^12^.

**Figure 2.**
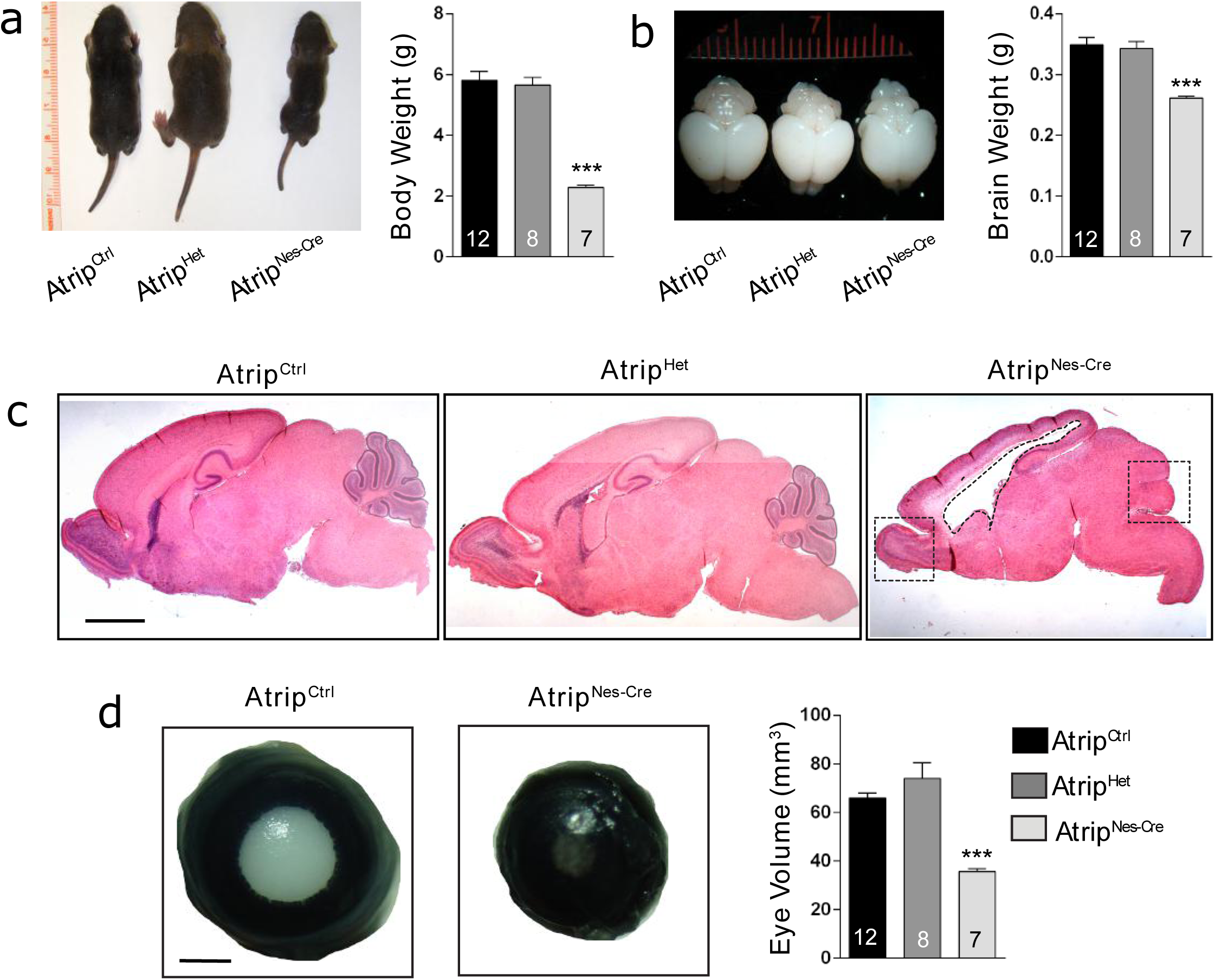
*Atrip* is essential for eye and CNS development: **(a,b)** Representative images of the body (a) and brain (b) and quantification of their weights in *Atrip*^*Ctrl*^, *Atrip*^*Het*^ and *Atrip*^*Nes-Cre*^ mice at P7. **(c)** Illustrative images of H&E staining of P7 brain sections from *Atrip*^*Ctrl*^, *Atrip*^*Het*^ and *Atrip*^*Nes-Cre*^ mice. The dotted line highlights the developmental defects observed in the cerebellum, cortex, hippocampus and striatum after *Atrip* inactivation. **(d)** Representative images and volume measurements of the eyes of *Atrip*^*Ctrl*^, *Atrip*^*Het*^ and *Atrip*^*Nes-Cre*^ mice at P7. Statistical analysis: one-way ANOVA followed by the Newman-Keuls posttest. The number of biological samples analyzed is indicated in the bars. Error bars indicate SEM; *** p<0.001.

### ATRIP is essential for lens development

To understand the function of ATRIP *in vivo*, we decided to focus our analysis on lens development. This tissue is a unique model for analyzing cell cycle dynamics *in vivo* because it is formed by only one progenitor cell type with well-characterized cell cycle progression. First, we evaluated the expression pattern of the *Atrip* and *Atr* genes in the mouse lens via real-time RT-PCR in various developmental stages. mRNA expression of both genes was found as early as embryonic day 12.5 (E12.5), and *Atr* and *Atrip* exhibited similar expression patterns (Supplementary figure 1a and b). To confirm the importance of *Atrip* for proper lens development, we generated *Atrip*^*Le-Cre*^ (*Atrip*^*Le-Cre*^ = *Atrip*^*F/F*^; *Le-Cre*^*+/-*^) mice, in which LoxP site recombination occurs in earlier stages of lens development (Ashery-Padan, 2000). ATRIP loss in the surface ectoderm (E9) led to aphakia (absence of lens) and eye growth defects (∼80% reduction in eye volume at P21) (Supplementary figure 1c and d). These findings reveal an essential function of *Atrip* for lens development.

### ATRIP loss induces cell death of lens progenitor cells

To investigate how ATRIP loss affects the maturation stages of lens development, we analyzed the morphology, cell proliferation and cell death of *Atrip*^*Nes-Cre*^ lenses. Nestin-Cre-mediated recombination occurs around E12.5-E13.5 in lens progenitor cells ^29^. At this developmental stage, the lens anterior epithelium is composed of cells that proliferate and migrate towards the equatorial region, where they exit the cell cycle and differentiate to form fiber cells ^30, 31^ (Figure 3a). Given the role of the ATRIP-ATR complex in the DNA damage response associated with DNA replication, we first asked whether *Atrip* deficiency induces the accumulation of DNA damage. At E17.5, the quantification of γH2AX-positive (γH2AX+) cells specifically in the progenitor cells of the lens epithelia revealed a marked ∼45-fold increase in the proportion of γH2AX+ cells in *Atrip*-deficient progenitor cells (Figure 3b). ATRIP loss also dramatically increased cell death, as revealed by the presence of TUNEL+ cells in the *Atrip*^*Nes-Cre*^ lens at E17.5 (Figure 3c). Consistent with apoptotic cell death, a ∼30-fold increase in the proportion of cleaved caspase-3 (cCasp3)-positive cells was observed in *Atrip*-deficient lens epithelia (Figure 3d).

**Figure 3.**
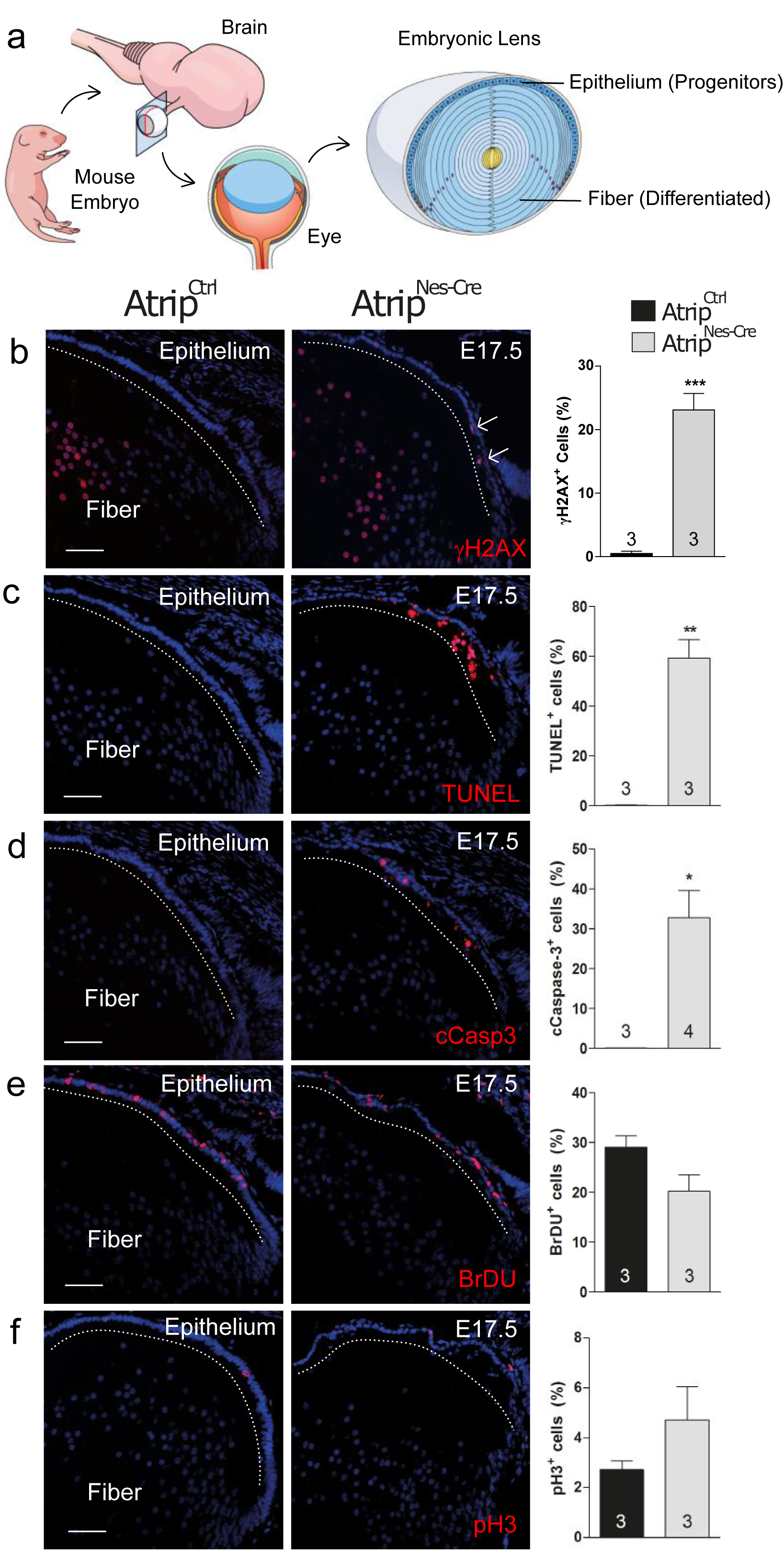
Inactivation of *Atrip* leads to DNA damage response and cell death in lens progenitor cells. **(A)** Schematic representation of the developing murine lens and its cell populations. Representative images and quantifications of E17.5 eye sections from control (*Atrip*^*Ctrl*^) and Atrip-deficient (*Atrip*^*Nes-Cre*^) mice stained for γH2AX **(b)**, TUNEL **(c)**, cleaved caspase-3 (cCasp3) **(d)**, BrdU **(e)** and pH3 **(f)** are shown. Arrows indicate positive cells among progenitor cells (lens epithelium). Note that both *Atrip*^*Ctrl*^ and *Atrip*^*Nes-Cre*^ lens fiber cells are γH2AX+. This staining profile is a classic hallmark of lens differentiation (Cvkel and Ashery-Padan, 2014; Cavalheiro et al., 2017). The number of biological samples analyzed is indicated in the bars. Statistical analysis: Student’s t-test. * p<0.05 ** p<0.01. *** p<0.001. Scale bar: 100 μm.

To test whether the proliferation rate of lens progenitor cells is affected by ATRIP loss *in vivo*, we scored the proportion of S-phase cells (BrdU+) in *Atrip*^*Nes-Cre*^ and *Atrip*^*Ctrl*^ lenses. No difference in the number of BrdU-incorporating cells was observed at E17.5 (Figure 3e). In addition, no change in the proportion of mitotic cells, as revealed by phospho-histone H3 (pH3) staining, was observed in *Atrip*-deficient progenitors (Figure 3f). These findings suggest that the ATRIP-ATR signaling pathway is key to preventing DNA damage accumulation and apoptosis in progenitor cells.

### Cell death induced by *Atrip* inactivation is TP53 dependent

DNA damage induces the stabilization of TRP53 and TRP53-mediated cell death (Meek, 2009). We asked whether the observed accumulation of DNA damage in *Atrip*-deficient lens progenitor cells leads to TRP53 stabilization and/or TRP53-mediated cell death. Immunostaining for total TRP53 revealed its stabilization in *Atrip*-deficient lens progenitor cells (Figure 4a). The phosphorylation of TRP53 (human S15/mouse S18) by PI3K-like kinases such as ATM and ATR is a well-characterized mechanism of TRP53 stabilization ^32, 33^. Consistent with the absence of TRP53 staining in control progenitor cells and with a phosphorylation-dependent stabilization of TRP53, phospho-TRP53+ progenitor cells were exclusively found in the *Atrip*^*Nes-Cre*^ lens (Figure 4b).

**Figure 4.**
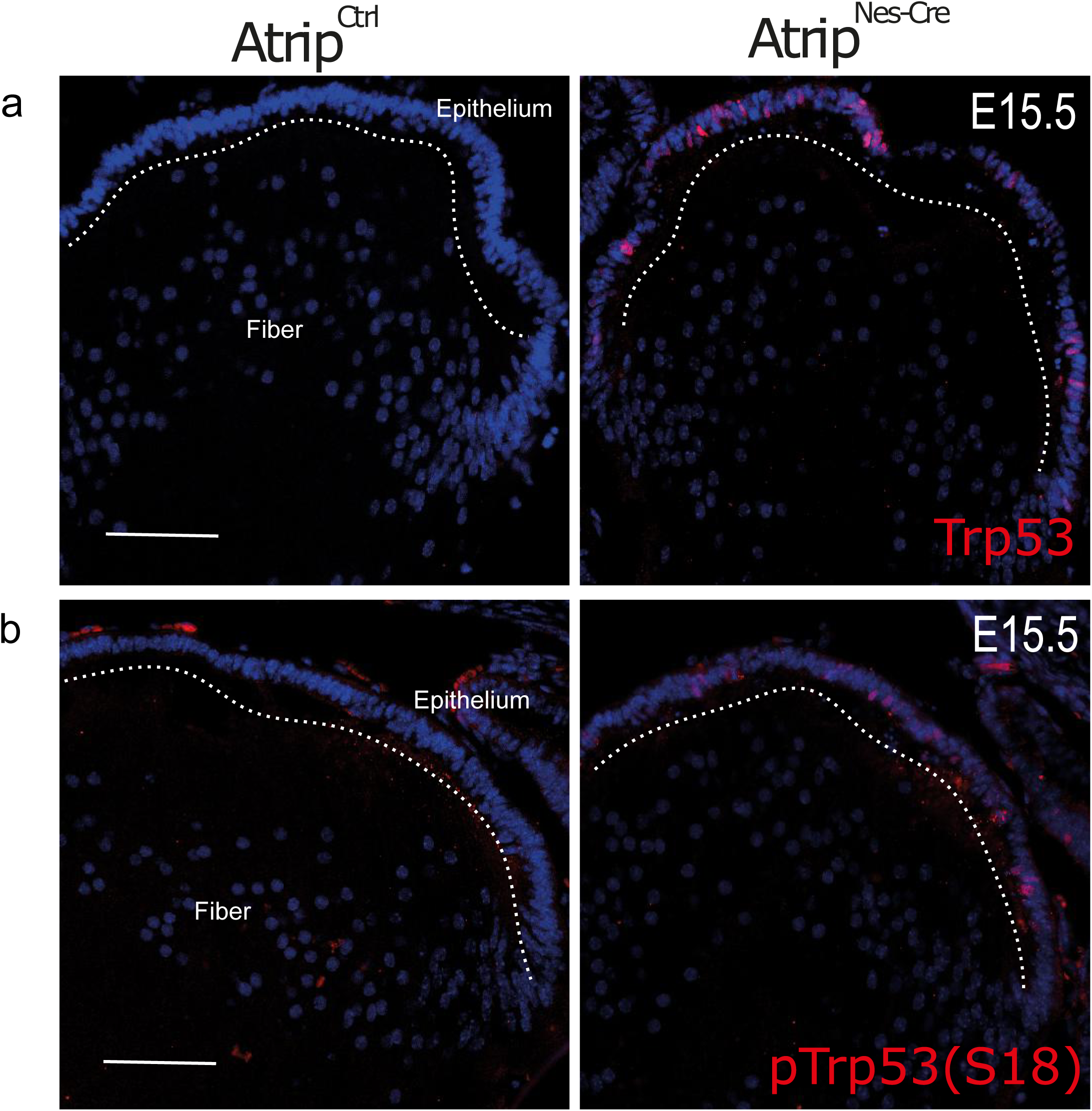
Inactivation of *Atrip* induced TRP53 stabilization in lens progenitor cells. Representative images of E15.5 lens sections from control (*Atrip*^*Ctrl*^) and Atrip-deficient (*Atrip*^*Nes-Cre*^) mice stained for Trp53 **(a)** and phospho-Ser18 Trp53 **(b)**. Scale bar: 100 μm.

In a number of cell types and species, TRP53 stabilization following DNA damage often results in TRP53-dependent cell cycle arrest and/or apoptosis (Meek, 2009). To test whether TRP53 mediates cell death following *Atrip* loss, we generated double-deficient *Atrip/Trp53* mice (*Atrip*^*Flox/Flox*^; *Trp53*^*Flox/Flox*^; *Nestin-Cre*^*+/-*^, referred to as *Atrip; Trp53*^*Nes-Cre*^ or cDKO). First, we evaluated whether *Trp53* inactivation affects cell cycle progression after *Atrip* inactivation. At E15.5, the quantification of S-phase (BrdU+) and mitotic cells (pH3+) revealed no difference between *Atrip*^*Nes-Cre*^ and *Atrip; Trp53*^*Nes-Cre*^ mice (Figure 5a and b), suggesting that TRP53 does not alter cell cycle progression following ATRIP loss. In contrast, as verified by the TUNEL assay (Figure 5c) and cCasp3 staining (Figure 5d), the inactivation of *Trp53* prevented the cell death induced by ATRIP loss in the E15.5 lens. We conclude that the apoptosis observed in embryonic *Atrip*-deficient lens progenitor cells is TRP53 dependent.

**Figure 5:**
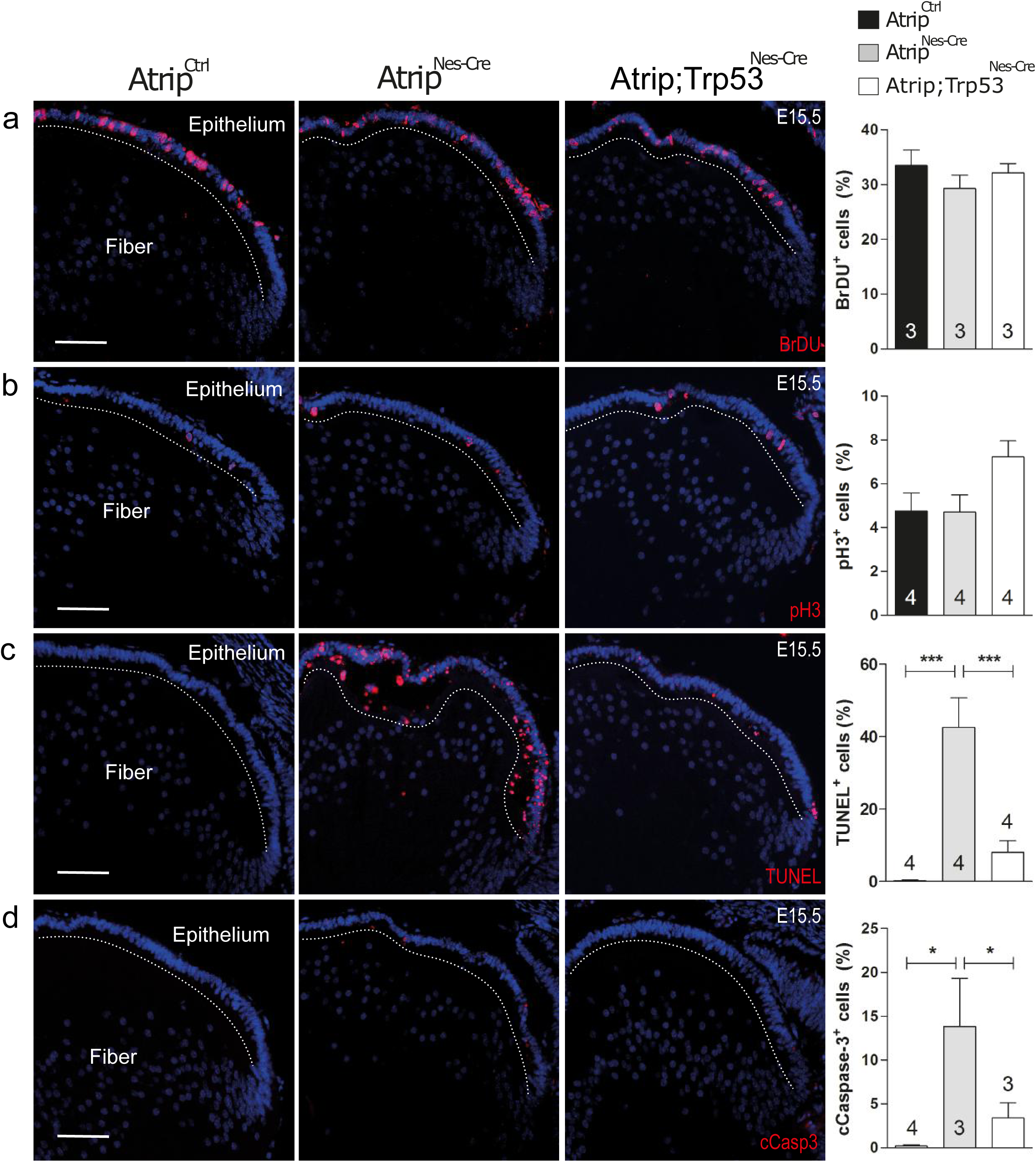
ATRIP loss leads to TRP53-mediated cell death in embryonic lens progenitor cells. Representative images of E15.5 eye sections from control (*Atrip*^*Ctrl*^), *Atrip*-deficient (*Atrip*^*Nes-Cre*^) and *Atrip* and *Trp53*-deficient (*Atrip; Trp53*^*Nes-Cre*^) mice stained for BrDU **(a)**, pH3 **(b)**, TUNEL **(c)** and cleaved caspase 3 (cCasp3) **(d)**. Statistical analysis: one-way ANOVA followed by the Newman-Keuls posttest. The number of biological samples analyzed is indicated in the bars. Error bars indicate SEM; * p<0.05. *** p<0.001. Scale bar: 100 μm.

### Simultaneous inactivation of *Atrip* and *Trp53* induces mitotic defects

To test whether the simultaneous inactivation of *Atrip* and *Trp53* affects the DDR in progenitor cells, we scored the proportion of γH2AX+ cells. Interestingly, even though *Trp53* inactivation abrogated the cell death induced by ATRIP loss, the proportion of γH2AX + progenitor cells was similar in *Atrip*^*Nes-Cre*^ and *Atrip; Trp53*^*Nes-Cre*^ mice (E15.5) (Figure 6a). This finding suggests that DDR activation is not altered after the inactivation of both *Atrip* and *Trp53*.

**Figure 6:**
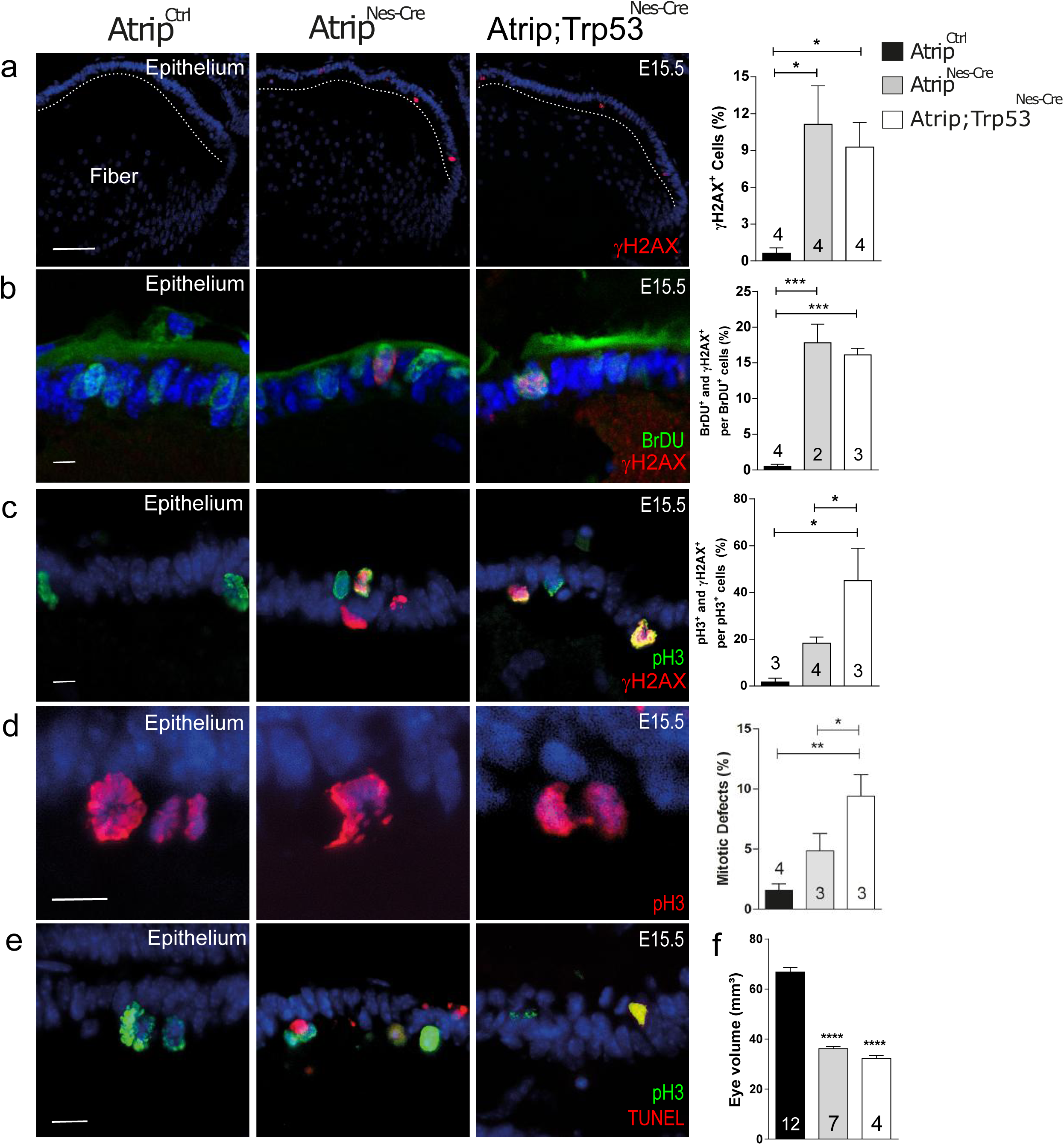
*Trp53* inactivation potentiates mitotic defects induced by ATRIP loss. Representative images and quantifications for the control (*Atrip*^*Ctrl*^), *Atrip*-deficient (*Atrip*^*Nes-Cre*^) and *Atrip* and *Trp53*-deficient (*Atrip;Trp53*^*Nes-Cre*^) lenses for γH2AX **(a)**, BrdU/γH2AX **(b)**, pH3/γH2AX **(c)**, mitotic defects (pH3) **(d)** and pH3/TUNEL double staining **(e)**. Apoptotic cells were observed during mitosis in *Atrip*^*Nes-Cre*^ and *Atrip;Trp53*^*Nes-Cre*^ samples (pH3/TUNEL double positive). **(f)** Eye volume quantification in control (*Atrip*^*Ctrl*^), *Atrip*-deficient (*Atrip*^*Nes-Cre*^) and *Atrip* and *Trp53*-deficient (*Atrip;Trp53*^*Nes-Cre*^) mice at P7. Statistical analysis: one-way ANOVA followed by the Newman-Keuls posttest. The number of biological samples analyzed is indicated in the bars. Error bars indicate SEM; * p<0.05 ** p<0.01. *** p<0.001. Scale bar: 100 μm for **(a)** and 10 μm for **(b, c, d** and **e**).

The ATRIP-ATR signaling pathway regulates faithful DNA replication, and its misregulation leads to replicative stress ^4^. Next, we examined whether ATRIP loss in the lens progenitor cells would increase DDR activation during S-phase by performing γH2AX and BrdU double staining. Consistent with the idea that *Atrip* inactivation increases replicative stress in lens progenitor cells, ∼17% of the BrdU-incorporating cells presented γH2AX staining after ATRIP loss, while only ∼0.5% were double positive in *Atrip*^*Ctrl*^. Moreover, cDKO and cKO lenses presented similar proportion of γH2AX and BrdU double-positive cells, showing that TP53 loss did not affect DDR activation following ATRIP loss in replicating lens cells (Figure 6b).

Studies have shown that *ATR* inactivation leads to the accumulation of mitotic defects in transformed cell lines ^4, 34^. To determine whether ATRIP loss induces DNA damage accumulation in lens mitotic cells, we evaluated E15.5 lenses double stained with γH2AX and pH3. ATRIP loss induced a 12-fold increase in the proportion of γH2AX+ cells within mitotic cells. Strikingly, while ∼18% of mitotic cells were γH2AX+ in *Atrip*^*Nes-Cre*^ mice, in the cDKO lens, we observed a further increase (∼45%) in mitotic cells accumulating DNA damage at E15.5. These findings indicate that the concomitant inactivation of *Trp53* and *Atrip* increased the accumulation of DNA damage in mitotic lens cells (Figure 6c).

Finally, we tested whether the increase in DDR activation is correlated with the accumulation of mitotic defects (bridges, multiple spindle pole formation and fragmented chromosomes). In the control lens, only ∼1.5% of mitotic cells presented defects, while ∼5% of mitotic cells exhibited defects following *Atrip* inactivation. Remarkably, a much higher proportion of mitotic cells (∼15%) presented defects in the cDKO lens epithelium (Figure 6d). Altogether, these findings suggest that in the absence of TRP53, some *Atrip*-deficient lens progenitor cells may escape the S/M checkpoint, enter mitosis, accumulate more DNA damage and eventually die during mitosis. Accordingly, double pH3 and TUNEL staining revealed apoptotic cells during mitosis in the cDKO lens at E15.5 (Figure 6e). Interestingly, although the concomitant inactivation of *Atrip* and *Trp53* reverted apoptosis induced by ATRIP loss, the eye volume of cKO and cDKO mice was similar at P7. This finding demonstrates that *Trp53* inactivation is not sufficient to rescue the eye growth impairment caused by ATRIP deficiency (Figure 6f). We conclude that, although TRP53 is able to rescue cell death, double inactivation of *Atrip* and *Trp53* increases mitotic defects and does not rescue eye growth defects observed after *Atrip* inactivation.

## Discussion

Our data demonstrate that ATRIP is essential for CNS and eye development. We showed that ATRIP protected progenitor cells from endogenous DNA damage. We found that a few days after Cre-mediated *Atrip* inactivation (∼3 days, at E15.5), lens progenitor cells underwent TRP53-dependent cell death during S-phase. Although *Trp53* inactivation rescued the apoptosis induced by ATRIP loss, DNA damage signaling was similar in *Atrip*-deficient and cDKO lens progenitor cells. In the absence of *Trp53, Atrip*-deficient lens progenitor cells survived through S-phase and showed increased accumulation of DNA damage and chromosomal defects in mitotic cells. Few of these cDKO progenitor cells died during mitosis, most likely due to the increased accumulation of DNA damage and, possibly, mitotic crisis. Accordingly, even though *Trp53* inactivation rescued the apoptosis caused by ATRIP loss, eye growth impairment was not reversed in cDKO mice. Taken together, our data demonstrate that the ATRIP-ATR complex is essential for protecting lens progenitor cells against endogenous DNA damage (Figure 7).

**Figure 7:**
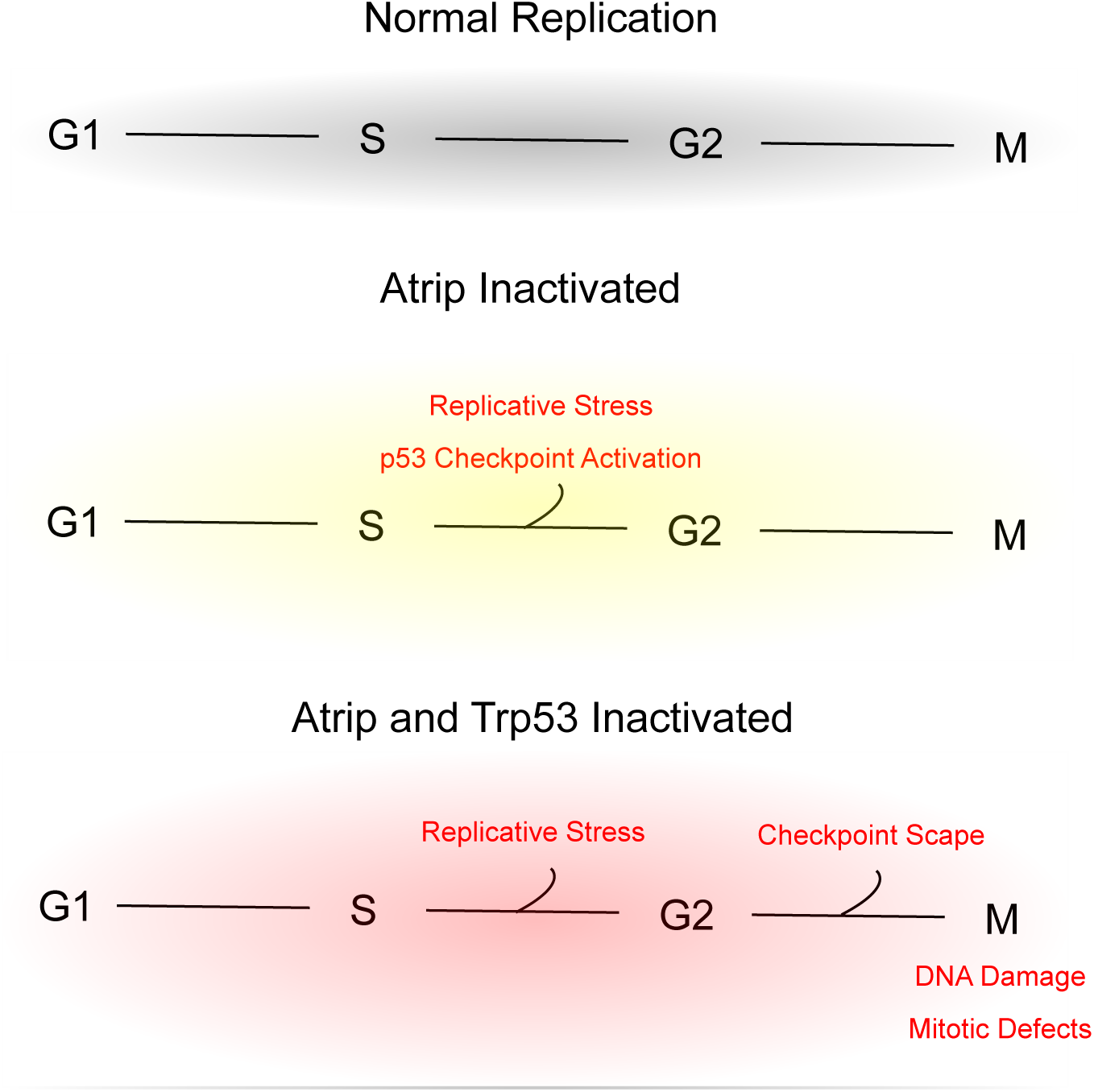
Effects of *Atrip* or *Atrip* and *Trp53* inactivation in lens progenitor cells.

Tissue growth defects are one of the main characteristics of Seckel syndrome patients, including facial dysmorphisms, microcephaly and ophthalmological defects ^35, 36, 37, 38^. Prior to our study, there was a lack of data supporting the idea that ATRIP could play a role during development. Here, we report the first mouse model of ATRIP loss of function. *Atrip* inactivation in the CNS and eye led to developmental growth impairment, including microcephaly, mirroring those associated with ATR loss ^12^. Moreover, *Atrip* inactivation decreased the ATR protein content, establishing a direct connection between ATRIP and ATR function *in vivo*. In summary, the role of ATRIP during development supports its relationship with Seckel syndrome etiology.

In the absence of ATRIP function, we observed increased activation of the DDR, as revealed by γH2AX and phospho-TRP53 staining. Following DSB formation, both ATM and DNA-PK are able to phosphorylate these targets ^32, 39, 40^. Although we did not determine which kinase was activated following ATRIP loss, our findings are consistent with the formation of DSBs and activation of ATM or DNA-PK or both. Nevertheless, it was shown that even of absence of the three main DDR kinases (ATM, ATR and DNA-PK), γH2AX phosphorylation and TRP53 activation are still detected, therefore we cannot exclude alternative pathways ^41^.

The depletion of the ATRIP-ATR complex was shown to increase underreplicated DNA, causing the accumulation of mitotic defects ^4, 42^ therefore, it is possible that the presence of underreplicated DNA during S-phase leads to the mitotic defects observed in *Atrip*-deficient lens progenitor cells. In this scenario, *Atrip* and *Trp53* double inactivation would increase the number of cells that are able to reach mitosis with underreplicated DNA, inducing a further increase in progenitors with mitotic defects. However, we cannot exclude the possibility of mitotic-specific roles of the ATRIP-ATR complex and TRP53. In fact, recent findings indicate that ATR is important for proper chromosomal segregation in cell lines ^5, 34^. Further studies are necessary to investigate the cell cycle phase-specific contributions of ATRIP -ATR and TRP53 during unchallenged replication *in vivo*.

*Atr* inactivation compromises the development of the cerebellum, which is associated with the accumulation of mitotic defects ^12, 19^. Interestingly, in contrast to what we observed in the developing lens, *Trp53* inactivation did not rescue the cell death or the increase in mitotic DNA damage induced by ATR loss in cerebellar progenitor cells ^19^. On the other hand, it was shown that *TP53*-deficient chronic lymphocytic leukemia (CLL) cells displayed an increase in mitotic defects after ATR inhibition ^43^. In addition, the accumulation of mitotic defects following *ATR* inactivation was also shown in transformed cell lines ^44^. These data indicate that ATRIP-ATR inactivation may have different outcomes depending on the cell type and genetic background. Since the ATRIP-ATR pathway is activated by replicative stress, we hypothesize that the levels of endogenous replicative stress may be at least partly responsible for these cell type-specific effects.

Currently, different ATR inhibitors are being successfully implemented in preclinical and clinical trials as monotherapy or in combination with other DNA-damaging drugs ^45, 46^, highlighting the relevance of studying the roles of the ATRIP-ATR complex *in vivo.* Our data reveal a novel contribution of ATRIP to the avoidance of DNA damage during development and provide a clear picture of the physiological consequences of inhibiting the ATRIP-ATR complex on *Trp53* proficient or deficient progenitor cells. The results presented here contribute to the understanding that ATR inhibitors may be toxic to cells with inactivated TP53, what may have important clinical implications.

## Acknowledgements

We thank Josee Guirouilh-Barbat, Emmanuelle Martini, Stephane Koundrioukoff, Leonardo K. Teixeira, Maurício Rocha-Martins and Ales Cvekl for the careful reading and discussion of the manuscript. We thank Isabele Pio Menezes and Severino Gomes for technical assistance and Dr. Graziela Ventura for assistance in image acquisition at the confocal microscopy facility of the Instituto de Ciências Biomédicas (ICB, UFRJ). This work was supported by the Brazilian National Council of Scientific and Technological Development (CNPq) (425556/2016-6, 439031/2018-4 and 313064/2017-2 to RAPM); FAPERJ (E-26/201.562/2014 and E-26/210.500/2019 to RAPM). This work was supported by the Deutsche Forschungsgemeinschaft (DFG): FR 2704/1-2 to POF for Initiation of International Collaboration. PG was the recipient of the PET PostDoc Fellowship from “Peter und Traudl Engelhorn Stiftung zur förderung der Biotechnologie und Gentechnik”.

## Supplementary figure legends

**Supplementary figure 1: *Atrip* and *Atr* expression pattern during lens development and inactivation of *Atrip* in the lens surface ectoderm. (a, b)** Real-time RT-PCR using specific primers for *Atr* and *Atrip* at 6 stages (E15.5, P0, P4, P9, P15. P60) of mouse lens development. *GPI-1* and *Actb* were used as internal control targets (n=3). **(c)** Representative images of P21 control *Atrip*^*Ctrl*^ and *Atrip*^*Le-Cre*^ eyes**. (d)** Eye volume measurements revealed severe impairment of eye growth after ATRIP loss in the lens surface ectoderm. The number of biological samples analyzed is indicated in the bars. Error bars indicate SEM. *** p<0.001.

