## Supplementary figures and images for "ATRIP protects progenitor cells against DNA damage *in vivo*"

### Sup. figure 1

Supplementary Figure 1

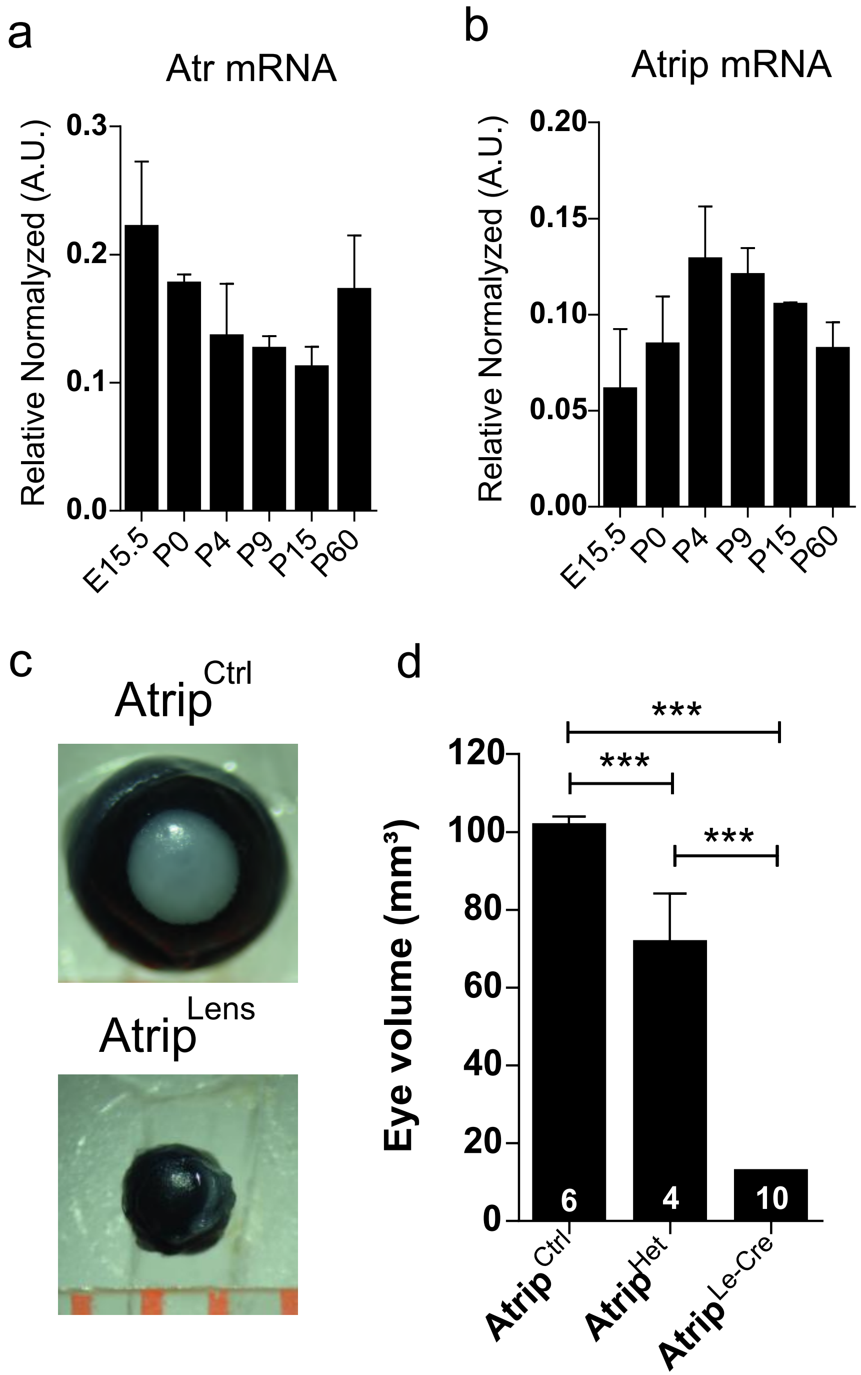


Matos-Rodrigues et al.
